# Bsp1, a fungal CPI motif protein, regulates actin filament capping in endocytosis and cytokinesis

**DOI:** 10.1101/2023.10.09.561521

**Authors:** Daniel R. Hummel, Markku Hakala, Christopher P. Toret, Marko Kaksonen

## Abstract

The capping of barbed filament ends is a fundamental mechanism for actin regulation. Capping protein controls filament growth and actin turnover in cells by binding to the barbed ends of the filaments with high affinity and slow off-rate. The interaction between Capping protein and actin is regulated by capping protein interaction (CPI) motif proteins. We identified a novel CPI motif protein, Bsp1, which is involved in cytokinesis and endocytosis in budding yeast. We demonstrate that Bsp1 is an actin binding protein with a high affinity for capping protein via its CPI motif. In cells, Bsp1 regulates capping protein at endocytic sites and is a major recruiter of capping protein to the cytokinetic actin ring. Lastly, we define Bsp1-related proteins as a distinct fungi-specific CPI protein group. Our results suggest that Bsp1 promotes actin filament capping by the capping protein. This study establishes Bsp1 as a new capping protein regulator and promising candidate to regulate actin networks in fungi.

## Introduction

In eukaryotes, a coordinated regulation of actin filament assembly and disassembly is required for diverse cellular processes, including cell migration, endocytosis and cytokinesis. A key regulator of actin assembly is the heterodimeric capping protein (CP) complex, which binds actin filament barbed ends and thereby prevents further barbed end growth (Isenberg et al., 1980; Carman et al., 2023). CP is necessary in maintaining the architecture of branched actin networks. The capping of filaments increases the polymerization rate of other filaments with free barbed ends (Carlier & Pantaloni, 1997; Pantaloni et al., 2001), enhancing the branched network assembly rates and thus actin-dependent forces that drive membrane deformation. In addition, association of CP to filaments inhibits actin disassembly and recycling. Thus, CP controls filamentous actin (F-actin) locally by defining filament lengths and globally by tuning actin turnover. In yeast, CP has a role in regulating actin polymerization at endocytic sites and thereby the growth of endocytic membrane invagination (Amatruda et al., 1992; Kim et al., 2004; Kaksonen et al., 2005). Furthermore, a recent study showed that CP also localizes to contractile actin ring in yeast, suggesting an additional role of CP in cytokinesis (Wirshing et al., 2023).

The activity of CP is finely tuned in cells (Edwards et al., 2014; Lappalainen et al., 2022). The uncapping of filament barbed ends is promoted by actin depolymerizing factor homology (ADF) domain proteins cofilin (Miyoshi et al., 2006; Wioland et al., 2017) and twinfilin (Hakala et al., 2021; Mwangangi et al., 2021). In animals, V-1 binds to CP and prevents its interaction with F-actin barbed ends through direct competition mechanism (Bhattacharya et al., 2006; Fujiwara et al., 2014). The activity of CP is regulated by CP interaction motif-containing proteins. The CP interaction (CPI) motif binds to CP and has an allosteric effect on the CP complex, leading to dissociation of V-1 from capping protein and thus promoting F-actin barbed end capping (Takeda et al., 2010; Fujiwara et al., 2014). The binding of CPI motif to CP also changes CP dissociation rates from F-actin and CP affinity to actin barbed ends (Hernandez-Valladares et al., 2010; McConnel et al., 2020; Takeda et al., 2021). In mammals several protein families, including CARMIL, Fam21, twinfilin, Cin85/CD2AP, CapZIP, and CKIP, are known to contain CPI motifs (Hernandez-Valladares et al., 2010; Edwards et al., 2014; McConnel et al., 2020; Hakala et al., 2018; Johnston et al., 2018; Takeda et al., 2021). Whereas no V-1 homologs have been identified in yeasts, the budding yeast *Saccharomyces cerevisiae* has two established CPI motif proteins, Twf1 and Aim21 (Falck et al., 2004; Johnston et al., 2018; Shin et al., 2018; Lamb et al., 2021). Both proteins localize to endocytic actin patches in yeast, regulating actin turnover and the activity of CP (Goode et al., 1998; Palmgren et al., 2001; Falck et al., 2004; Johnston et al. 2015, Farrell et al., 2017; Shin et al., 2018; Lamb et al., 2021).

Here we identify Bsp1 as a third fungal CPI motif protein. We found that Bsp1 promotes the localization of CP to both actin patches and the contractile ring through direct interactions with both F-actin and CP. Our sequence analysis reveals that the CPI motif of Bsp1 is closer to those of CKIP and Fam21, pointing to a pro-capping function of Bsp1. Combined, our data suggests that Bsp1 is a pro-capping factor in endocytosis and cytokinesis.

## Results and Discussion

### Bsp1 domains and localization

Bsp1 localizes to endocytic actin patches and cytokinetic actin rings (Drees et al., 2001; Wright et al., 2008; Tonikian et al., 2009). Bsp1 with Iqg1 is essential for actin ring formation and the deletion of both genes together results in cell division defects (Wright et al., 2008). Based on AlphaFold structure predictions, yeast Bsp1 is a largely unstructured protein without any known domains. The N-terminal part of Bsp1 is proline-rich and therefore could potentially be bound by a diverse set of protein-protein interaction domains, e.g. SH3 domains, as well as by actin or profilin-bound actin (He et al., 2009; Tonikian et al., 2009; Feliciano et al., 2015; Bieling et al., 2018). We searched for short peptide motifs in the Bsp1 sequence and identified a region with two sequences that resembled WH2 domains, a F- and G-actin-binding domain family (Paunola et al., 2002; Barzik et al., 2005; Co et al., 2007; Breitspechter et al., 2008; Breitspechter et al., 2011; Carlier et al., 2013). Furthermore, we detected a putative CPI motif at the C-terminus of the protein. This motif shares similarities to CPI motifs of CARMILs and CKIPs, but differs from CPI motif of twinfilins (Fig. 1A). The putative WH2 and CPI sequences together with the localization of Bsp1 actin structures suggest a role in actin regulation.

**Figure 1:**
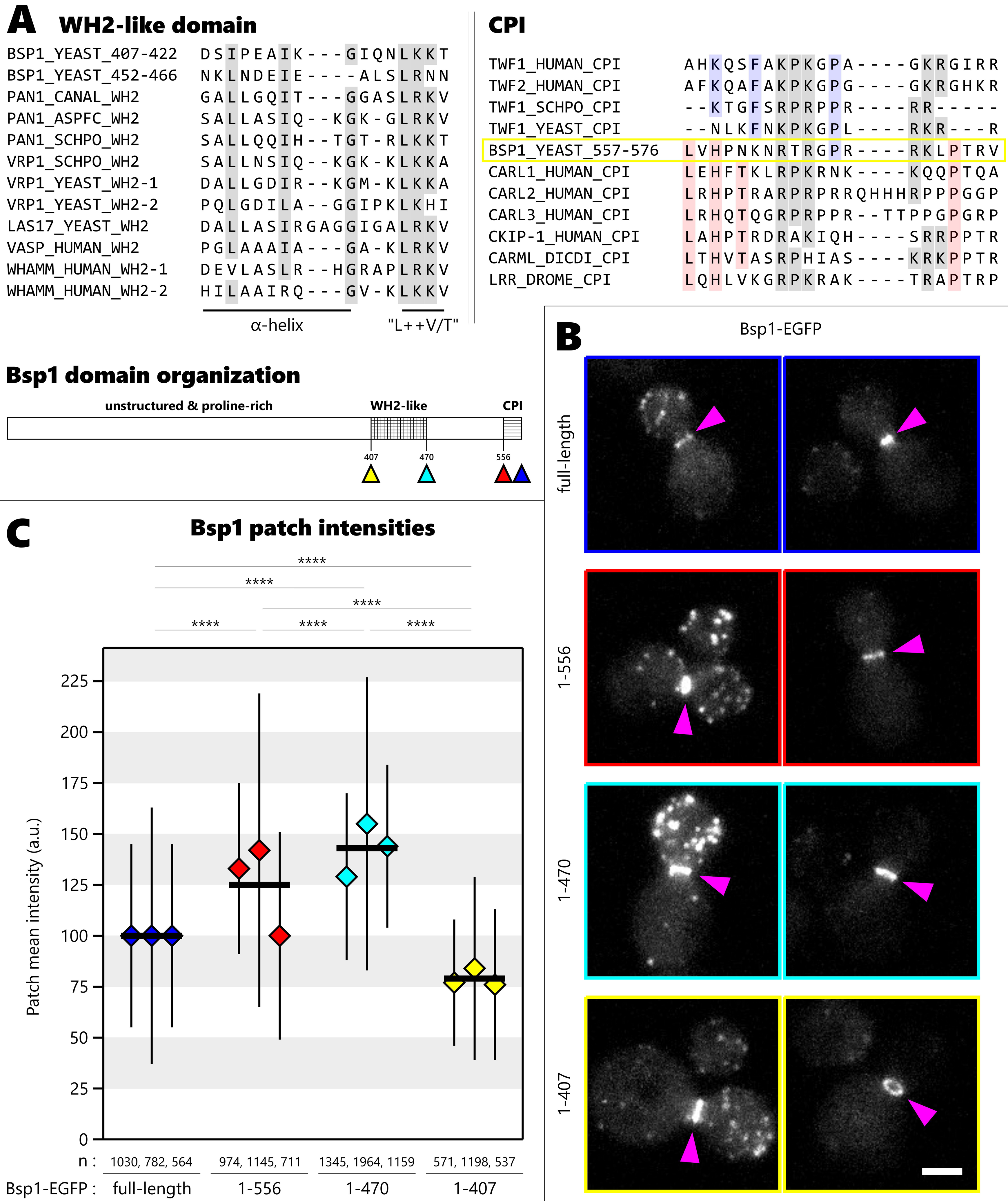
Bsp1’s WH2-like domain and CPI regulate Bsp1 recruitment to endocytic sites. (A) Top left, various WH2 domain sequences compared to Bsp1 sequences; key residues of WH2 domains are highlighted in gray. Top right, Bsp1 CPI sequence compared to twinfilin-like and CARMIL-like CPIs; key residues of twinfilin-like CPIs, of CARMIL-like CPIs and shared key motifs are highlighted in blue, red and gray, respectively. Bottom left, domain organization of Bsp1; colored arrows indicate position of truncations and GFP-tagging for B) and C). (B) Localization of Bsp1-EGFP, Bsp1^1-556^-EGFP, Bsp1^1-470^-EGFP and Bsp1^1-407^-EGFP. Pink arrows highlight cytokinetic ring structures. Images of representative cells are projections of z-stacks. Scale bar: 2 µm. (C) Mean intensity of patches with standard deviation. Thick black lines are the mean of means. Number of patches analyzed per experiment is indicated by n. Pairwise comparison based on Welch’s t-test (****, p < 0.0001).

To study the role of CPI motif and WH2-like domains in the Bsp1 recruitment to actin structures, we GFP-tagged Bsp1 C-terminally at its endogenous gene locus (Fig. 1A). We also designed three C-terminally GFP-tagged truncation mutants: Bsp1^1-556^, Bsp1^1-470^ and Bsp1^1-407^ (Fig. 1A). We observed a heterogeneous expression level of Bsp1 across the cell population that appeared to be independent of cell cycle state, which has not been reported for other actin patch proteins (Fig. S1A). Full-length Bsp1 localized to endocytic patches and the contractile ring consistent with previous findings (Fig. 1B) (Drees et al., 2001; Wright et al., 2008). Occasionally, we observed in some cells a filamentous localization of Bsp1 reminiscent of actin cables (Fig. S1B). Compared to full-length Bsp1, all Bsp1 mutants showed a similar overall localization to actin patches and contractile rings. However, more detailed analysis revealed that compared to the full-length protein, both Bsp1^1-556^ and Bsp1^1-470^ were more enriched at actin patches (fig. 1C). In contrast, the level of Bsp1^1-407^ mutant at patches was significantly decreased (fig. 1C). This suggests that the WH2-like domain promotes Bsp1 recruitment to actin patches and that the C-terminal region, including the CPI motif, negatively regulates Bsp1 recruitment to actin patches. Alternatively, these Bsp1 recruitment phenotypes may reflect changes in the amount of actin at endocytic sites caused by the Bsp1 mutations. None of our mutants showed complete mislocalization, indicating that the proline-rich N-terminal part of the protein also has a role in the localization of Bsp1.

### Bsp1 binds actin filaments and capping protein *in vitro*

Bsp1 localizes to different actin filament populations in yeast cells. Bsp1 might interact directly with F-actin through its putative WH2-like domains. To test this hypothesis, we purified untagged full length Bsp1 protein and tested its affinity to actin filaments with co-sedimentation assay. Using polymerized human platelet actin, we observed an increasing amount of Bsp1 protein in the pellet fraction after ultracentrifugation upon increased concentration of actin filament (Figure 2A-B). These results indicate that full length Bsp1 binds directly to F-actin.

**Figure 2.**
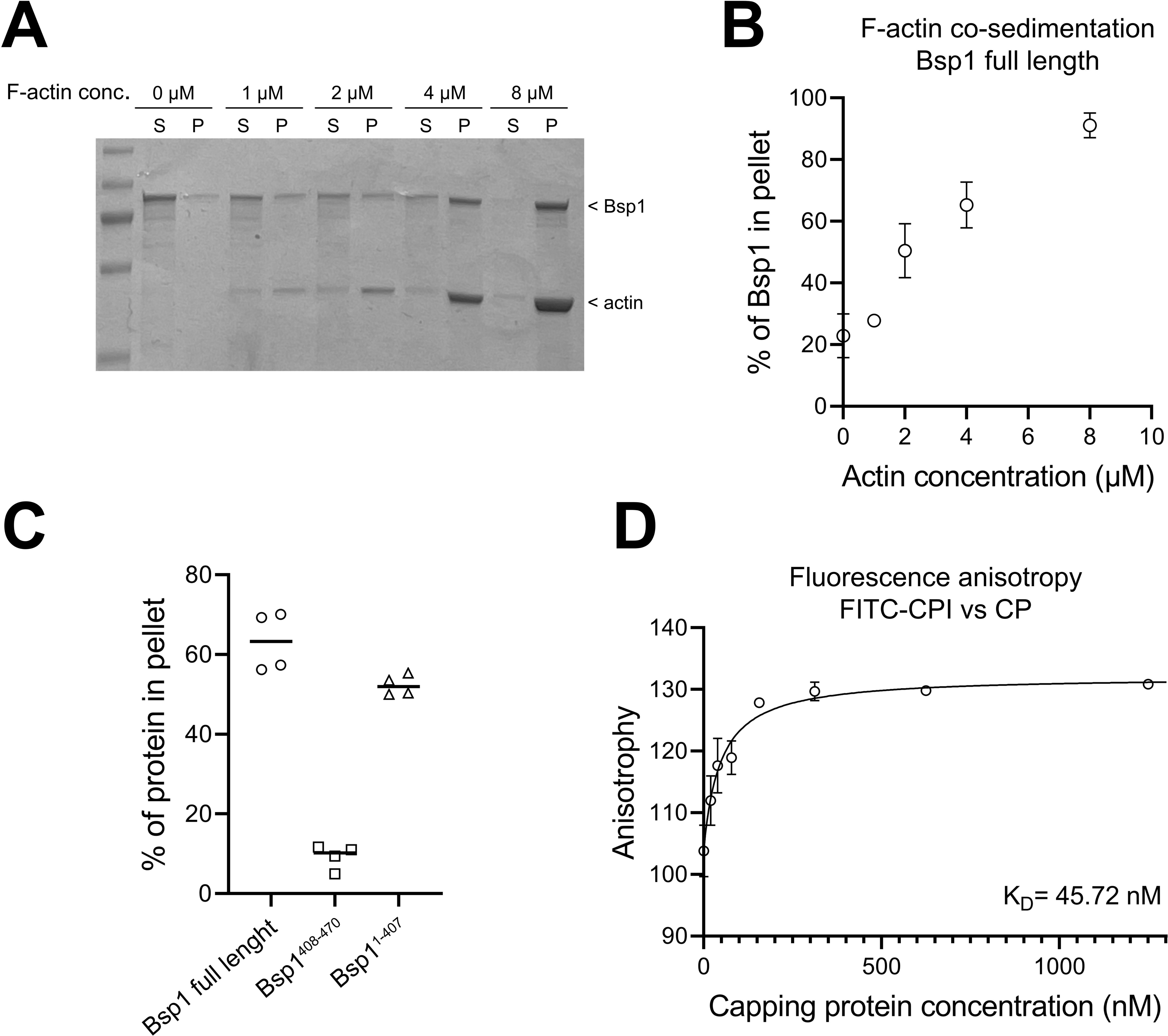
Bsp1 interacts with F-actin and CP in vitro. (A) A representative example of SDS-PAGE gels of F-actin co-sedimentation assay. 2 µM full length Bsp1 and indicated amount of F-actin were used. (B) F-actin co-sedimentation assay with 2 µM Bsp1 and indicated concentration of F-actin. Data represents a mean of three experiments with standard deviations shown. (C) F-actin co-sedimentation assay with 2 µM full length Bsp1, Bsp1^1-407^, and GST-Bsp1^408-470^ with 2 µM F-actin. All data points and means are shown. (D) Fluorescence anisotropy assay with 100 nM FITC-labeled CPI peptide and indicated concentration of yeast capping protein. Data represents a mean of three experiments with standard deviations shown. Data is fitted with a one-site binding equation (see materials and methods).

To test if the two putative WH2 domains facilitate the interaction of Bsp1 with actin filaments, we purified both a truncated Bsp1^1-407^ lacking the two WH2-like domains and the C-terminal, CPI motif, as well as the two WH2-like domains conjugated with GST (GST-Bsp1^408-470^). We observed strong interaction of full length Bsp1 and GST-Bsp1^408-470^ with actin filaments, whereas Bsp1^1-407^ did not interact with F-actin (Figure 2C).

Next we wanted to study if Bsp1 interacts directly with CP. To this end, we purified recombinant yeast CP and synthesized the CPI motif peptide of Bsp1 and labeled it with Cy5. By measuring the fluorescence anisotropy of labeled CPI motif peptide, we observed a strong affinity of Bsp1 CPI motif peptide to CP with the dissociation constant K_D_ being 45 nM (Fig 2D). Compared to other known CPI motifs (McConnell et al. 2020), Bsp1 binds CP with affinity similar to CIN85 and CD2AP, whereas mammalian CARMILs bind to CP with much stronger affinity. Notably, mammalian twinfilin has much lower affinity to CP compared to Bsp1 (Hakala et al., 2018; McConnell et al., 2020), suggesting that these proteins have a different effect on capped F-actin barbed ends. In summary, our *in vitro* binding assays show that Bsp1 has a strong affinity to CP with its C-terminal CPI motif, and to F-actin through the two WH2-like motifs.

### Bsp1 regulates actin patch composition

To further investigate the role of Bsp1 in endocytic actin regulation, we analyzed the intensities of GFP-tagged Cap1 (actin filament capping complex α subunit), Twf1 (twinfilin), Sac6 (fimbrin), Abp1 (Actin-binding protein 1) and Arc18 (Arp2/3 complex subunit) at the actin patches. Unlike Bsp1, these endogenously expressed proteins showed uniform expression levels across the population of cells (fig. S2). The peak intensities of Twf1, Cap1 and Abp1 patches were significantly reduced (19, 17 and 13 %, respectively) in *bsp1Δ* cells compared to WT cells (Fig. 3A). A reduced number of Cap1 molecules at endocytic sites in *bsp1Δ* cells suggests that Bsp1 either promotes actin filament capping or stabilizes CP at filament barbed ends. The similarity of the Cap1 and Twf1 phenotypes could be explained by Cap1-dependent recruitment of Twf1 (Palmgren et al., 2001). In contrast, Sac6-EGFP patch peak intensities were not significantly different between WT and *bsp1Δ* cells, suggesting that actin cross-linking is unaffected (Fig. 3A). Strikingly, the average peak patch intensity of Arc18-eGFP was increased by 17% in *bsp1Δ* cells over WT (Fig. 2A), suggesting that the endocytic actin network in *bsp1Δ* cells is more branched than in WT cells. Remarkably, a fluorescent phalloidin staining showed that the overall F-actin amount in endocytic patches was not altered in *bsp1Δ* compared to WT cells (Fig. 3B).

**Figure 3:**
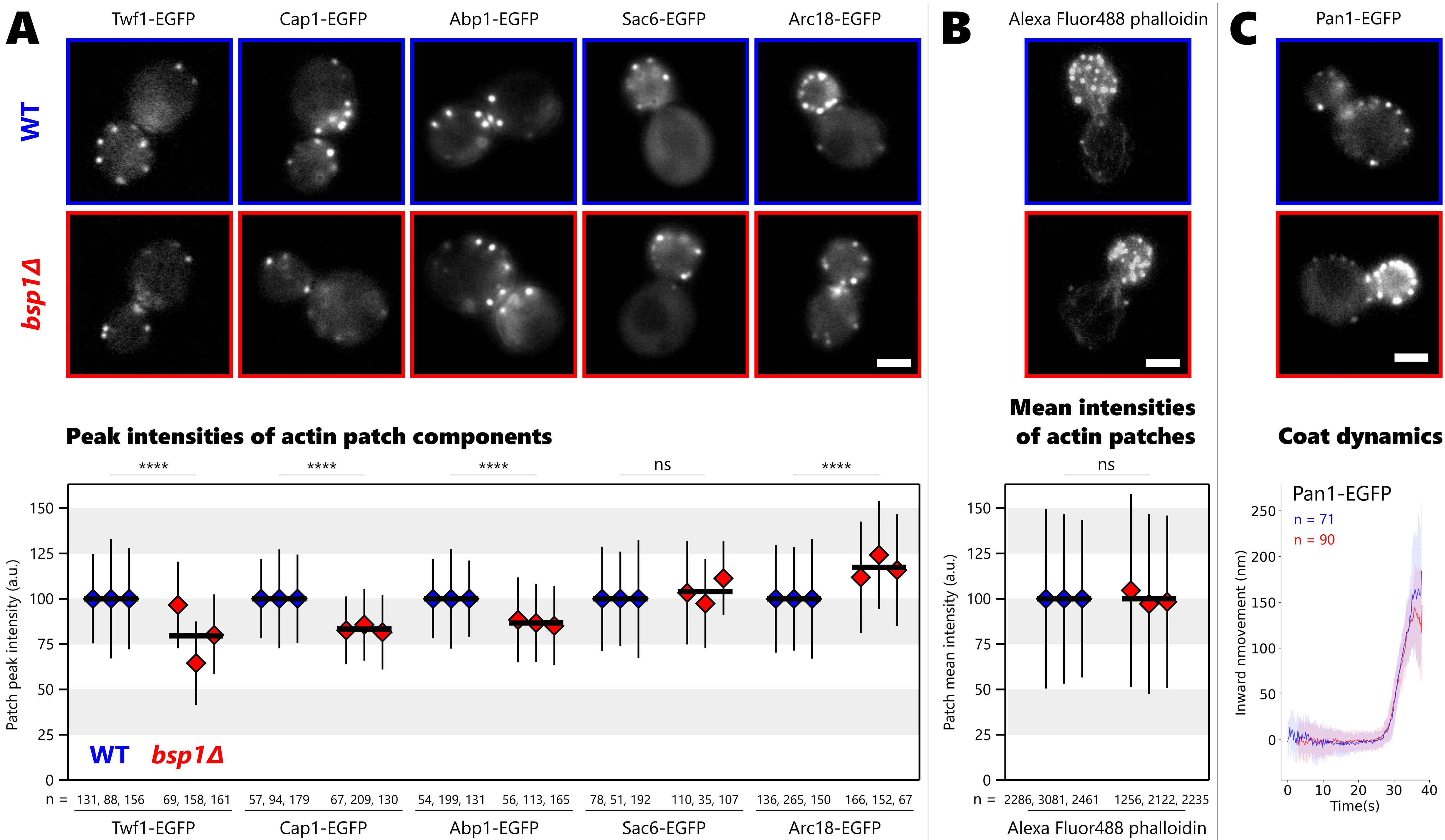
Bsp1 regulates actin patch composition at endocytic sites. (A) Localization of different actin patch components in WT (blue) and bsp1Δ cells (red). Images of representative cells are single frames from movies. Plot shows peak intensity of protein patches as mean with standard deviation. Thick black lines are the mean of means. Number of patches analyzed per experiment is indicated by n. Pairwise comparison based on Welch’s t-test (ns, not significant; ****, p < 0.0001). (B) Localization of phalloidin-stained actin patches in WT (blue) and bsp1Δ cells (red). Images of representative cells are projections of z-stacks. Plot shows average intensity of phalloidin patches as mean with standard deviation. Thick black lines are the mean of means. Number of patches analyzed per experiment is indicated by n. Pairwise comparison based on Welch’s t-test (ns, not significant). (C) Pan1-EGFP patches in WT (blue) and bsp1Δ cells (red). Images of representative cells are single frames from movies. Plot depicts median centroid movement (solid lines) with median absolute deviations (shaded areas). Images of representative cells are single frames from movies. Scale bars: 2 µm.

Taken together these results suggest that relative to WT, in *bsp1Δ* cells the endocytic patches have similar amounts of F-actin and cross-linkers, while Arp2/3-branching is increased and the amount of CP, Twf1 and Abp1 is decreased. Actin network organization is likely tuned differently in the mutants with less capped and more branched filaments. These findings are consistent with either a pro-capping or F-actin cap stabilizing role for Bsp1. In addition, Abp1 interacts with Cap1, Sac6 and the Arp2/3 complex, but the amount of Abp1 in the actin patch was reduced in *bsp1Δ* cells, similar to Cap1. This observation suggests that Abp1 recruitment follows CP, and not Arp2/3 or Sac6. In addition, Bsp1 proline rich regions might indirectly regulate the Arp2/3 complex, through other actin binding proteins via SH3 domain-interaction network. (Tong et al., 2002; Tonikian et al., 2009; Hummel and Kaksonen, 2023). As actin network composition in *bsp1Δ* cells differs from WT, the actin-dependent growth of the endocytic invagination might be affected. We used the motility of Pan1-EGFP, a coat associated protein, as a proxy for invagination growth (Kaksonen et al., 2003; Picco et al., 2015). The analysis of Pan1 motility showed that membrane invagination and vesicle formation are not affected by the deletion of Bsp1 (Fig. 3C). Hence, Bsp1-associated actin patch composition differences are mild enough so that endocytic invagination effects are not detectable. It is possible that actin regulation by Bsp1 becomes important for stress response or plays a role in balancing the competition for actin between cytokinetic and endocytic sites.

A deletion or mutation of the CPI motif protein Aim21 reduced Cap1 amounts at endocytic sites in previous studies similar to our Bsp1 mutants (Farrell et al., 2017; Lamb et al., 2021). In contrast, we found that Bsp1 loss decreased Abp1 numbers in actin patches, while the deletion of Aim21 increased Abp1 numbers in patches (Shin et al., 2018). Hence, the two CPI motif proteins Bsp1 and Aim21 have distinct functions. Bsp1 co-localizes with the endocytic actin network while Aim21 is co-localized with the WASP module proteins at the base of the endocytic invagination (Tonikian et al. 2009). These differences in localization of Bsp1 and Aim21 may correlate with their unique activities related to CP. Twf1, like Bsp1, is a component of the actin module, but the localization of Twf1 is dependent on its CP interaction (Palmgren et al., 2001). The CPI of Bsp1 is not necessary for its recruitment and may even hinder it. The distinct recruitment mechanisms suggest that these CPI proteins have different roles in CP regulation. Our data supports the idea that Bsp1 promotes barbed end capping by CP, whereas at least mammalian twinfilin-1 was shown to uncap filament barbed ends (Hakala et al., 2021; Mwangangi et al., 2021). While yeast and mammalian twinfilins likely have conserved biochemical functions, we note that to date yeast Twf1 has not been shown to uncap filament barbed ends.

### Bsp1 recruits CP to the bud neck

The actin patch at the endocytic sites is nucleated by the Arp2/3 complex, while the cytokinetic contractile actin ring is nucleated by formins (Pruyne et al., 2004). CK666 is an Arp2/3 inhibitor, which blocks actin patch formation in yeast without perturbing the contractile ring. A recent study found that Cap2 normally localizes to both the endocytic patches and the contractile ring, however, upon CK666-treatment, Cap2 localizes only to the contractile ring (Wirshing et al., 2023). We tested if the Bsp1 CPI motif could regulate recruitment of CP to the contractile ring. In agreement with the previous study (Wirshing et al., 2023), we observed Cap1-eGFP localization both to endocytic patches and the contractile ring. However, the bud neck area is rich in endocytic events, which makes quantification of Cap1 ring localization difficult. CK666-treatment made ring formation of Cap1-EGFP more apparent due to a strong reduction of Cap1 patches. Thus, we tested whether the Bsp1 CPI motif can regulate Cap1 recruitment to the bud neck in CK666-treated cells. We counted the number of Cap1-EGFP ring structures per field-of-view (FOV). Compared to WT levels, the number of cells with Cap1-EGFP localized to the cytokinetic ring was significantly decreased in both *bsp1Δ* cells and in *bsp1*^1-556^ cells (fig. 4A). Both Bsp1-EGFP and Bsp1^1-556^-EGFP localized to the contractile ring in a similar manner (Fig. 4B). This data shows that Bsp1 with its CPI motif is the major recruitment factor for CP and thus a key regulator of actin filament capping at the yeast contractile ring.

**Figure 4:**
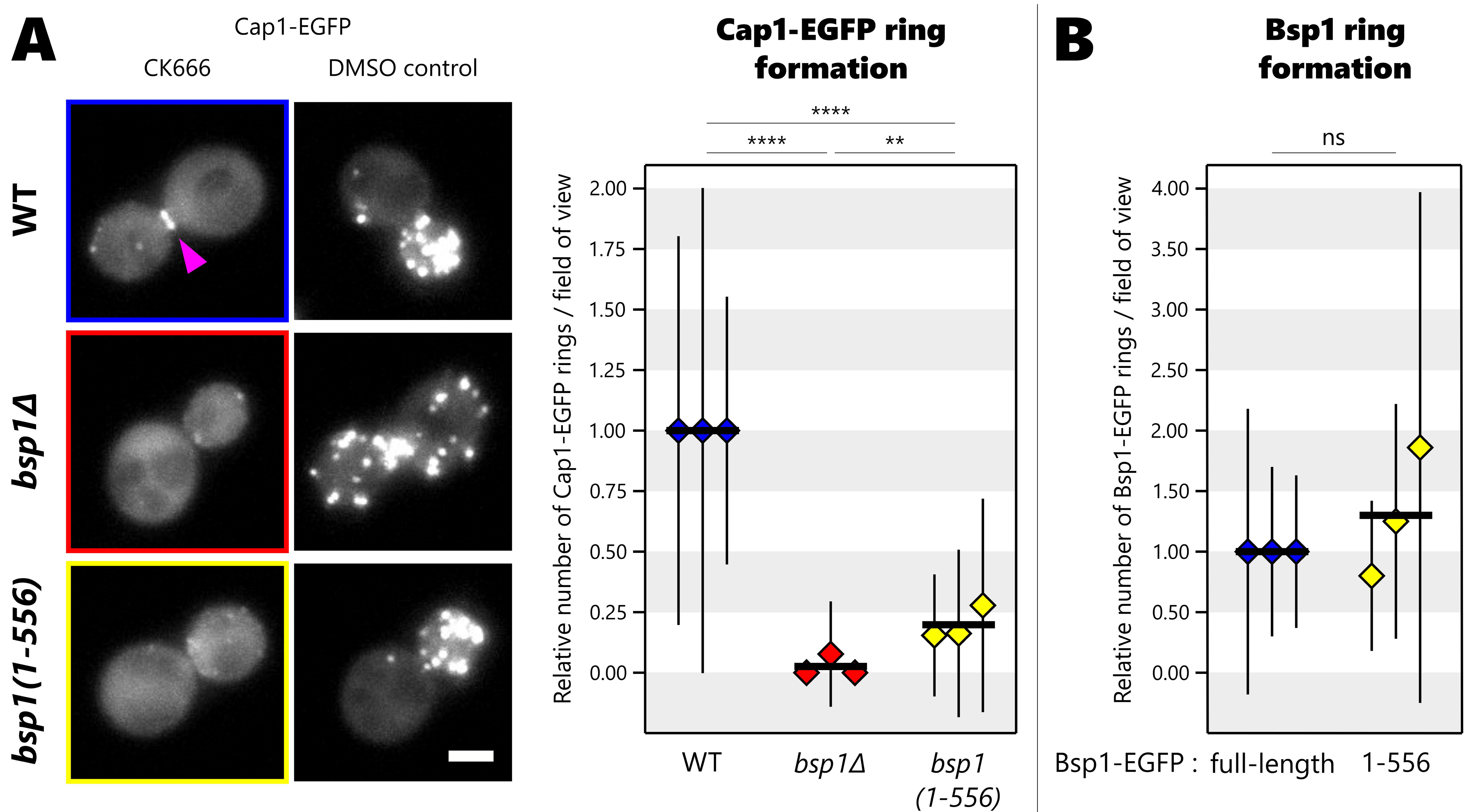
Bsp1 regulates Cap1 recruitment at contractile rings. (A) Cap1-EGFP localization in CK666-treated cells and DMSO control in WT (blue), bsp1Δ (red) and bsp1^1-556^ cells (yellow). Images of representative cells are projections of z-stacks. Pink arrows highlight cytokinetic ring structures. Plot shows mean number of cells with a Cap1-EGFP localizing to the cytokinetic ring in CK666 treated cells per field-of-view, error bars are standard deviation. Thick black lines are the mean of means. Pairwise comparison based on Welch’s t-test (**, p < 0.01; ****, p < 0.0001). Scale bars: 2 µm. (B) Bsp1-EGFP (blue) and Bsp1^1-556^-EGFP (yellow) localization in CK666-treated cells. Plot shows mean number of cells with a Bsp1 localizing to the cytokinetic ring in CK666 treated cells per field-of-view, error bars are standard deviation. Thick black lines are the mean of means. Pairwise comparison based on Welch’s t-test (ns, not significant).

The CP localization defect was milder in *bsp1*^1-556^ than in *bsp1Δ*, suggesting that Bsp1 may have several mechanisms of regulating CP recruitment. Moreover, CP recruitment to both endocytic sites and the contractile ring was reduced in *bsp1Δ*, suggesting increased levels of free, inactive cytoplasmic CP. We speculate that Bsp1 may have a CP-activating function analogous with mammalian CARMIL-like CPI motif proteins. Alternatively, as Bsp1 can bind actin *in vitro* (fig. 2), Bsp1 may stabilize barbed end-bound CP through simultaneous binding to both CP and actin filaments.

### Bsp1s are a CPI motif protein family of dikarya fungi

We examined the Bsp1 amino acid sequence in respect to other orthologs and known CPI motif proteins. With step-wise and reciprocal BLAST searches we were able to identify orthologs in most dikarya, a fungal clade that contains over 90% of known fungal species. We found three distinct groups of Bsp1 orthologs (fig. 5A). The orthologs in the first group, which we term Group I, have an N-terminus that resembles budding yeast Bsp1 in organization, but have up to three additional C-terminal ADF/gelsolin domains after the CPI motif. Both dikarya subgroups, the ascomycetes and basidiomycetes, contain Group I orthologs, which likely represent an ancestral Bsp1 organization. The second group is found in Saccharomycotina subphylum, which includes *S. cerevisiae* Bsp1. Group II proteins lack the ADF/gelsolin domains and terminate with a CPI motif. The WH2 domains were only detected in some Group II orthologs. The third group comprises proteins found in Ustilaginomycotina subphylum. These Bsp1 orthologues fused with the Ade8 gene, a purine biosynthesis enzyme. Ade8 (a GART domain), is inserted in the center of Group III Bsp1s and appears to have replaced the CPI motif (fig. 5A). This may hint at a unique feedback between ADP/ATP synthesis and actin, a major consumer of ATP.

**Figure 5.**
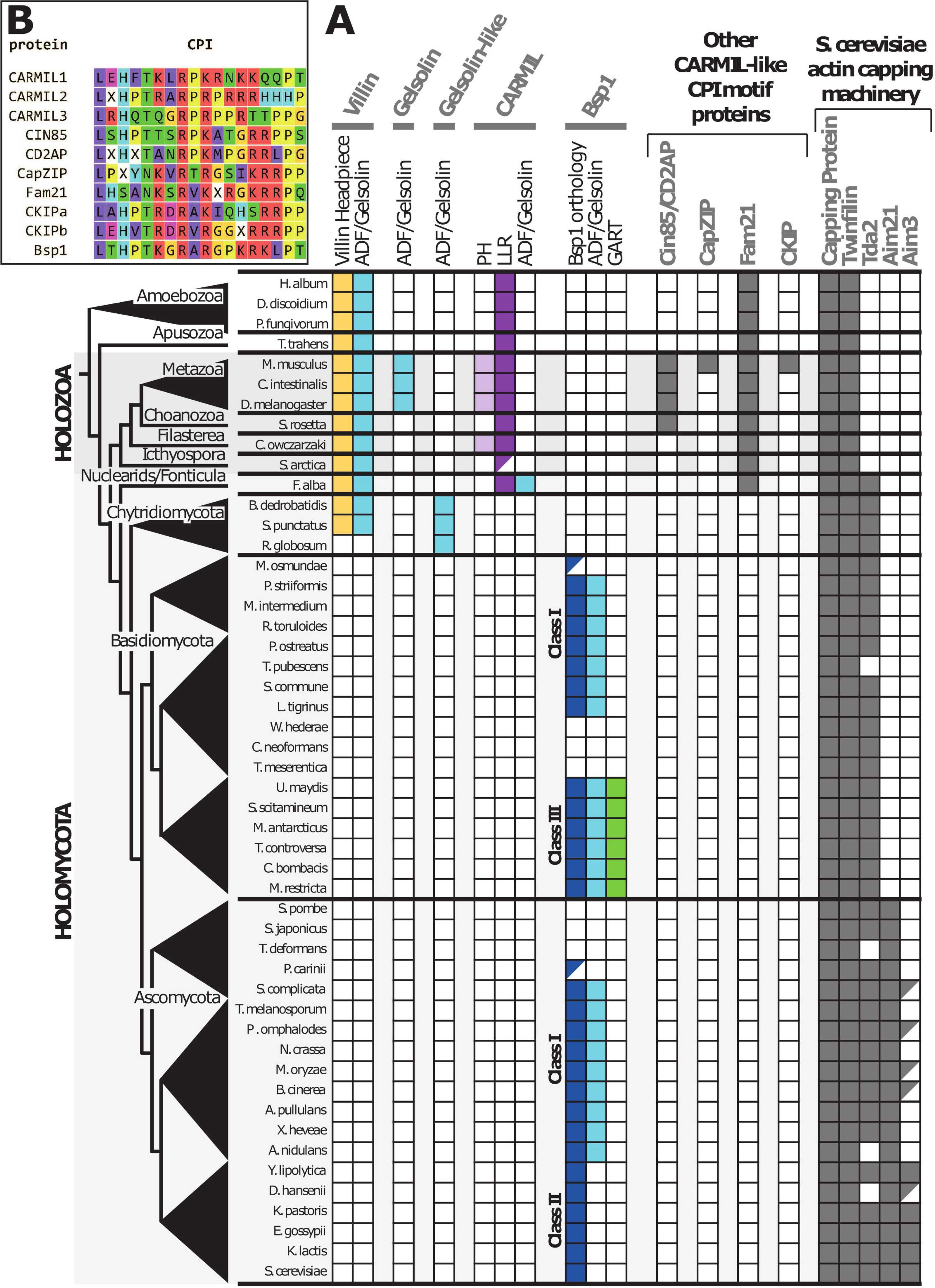
Bsp1 and CP regulation orthology. (A) Table lists the presence of at least one protein ortholog (top) in each species (left). Approximate clade location for each species on the tree of life (far left). For villin, gelsolin, gelsolin-like, CARMIL and Bsp1. Presence or absence of specifically indicated domains is shown (colors). Partially filled boxes indicate an orthology that is either truncated or incomplete in the database. (B) Bsp1 family CPI consensus sequence alignment with other CPI-containing protein family consensus sequences.

Many fungal Bsp1 proteins have been annotated as villin/gelsolin proteins due to the presence of C-terminal ADF/gelsolin domains. Canonical villins occur in diverse species including plants, amoeba and animals and contain several ADF/Gelsolin domains in succession, followed by a C-terminal villin headpiece domain (George et al., 2020; Huang et al., 2014). Proteins with this organization were not detected in dikarya species (fig. 5A). Gelsolins resemble villin ADF/gelsolin domain organization, but lack a headpiece and appear restricted to muscle-containing animal species (fig. 5A). A gelsolin-like protein occurs with villin in the fungal group Chytridiomycota, but it is unclear if this is a novel fungal convergence, a Bsp1 variant or has another origin (Fig. 5A). A connection between true villins/gelsolins and Bsp1s remains unclear, given their few ADF/Gelsolin domains, lack of a villin headpiece and CPI motif.

We sought to identify if Bsp1s were evolutionarily related to other CPI motif proteins. The CPI motif proteins Cin85/CD2AP, CKIP, and CapZIP were only detected in holozoans, while CARMILs and Fam21 proteins appeared more ancient (fig. 5A). Canonical CARMIL proteins contain an N-terminal PH domain, LLR domains and a CPI motif, however the PH domain has been lost in several lineages. Fungi were not found to contain CARMIL orthologs, however the holomycotan *Fonticula alba* contained a CARMIL that had uniquely added C-terminal ADF/gelsolin domains. The unstructured Fam21 proteins are a subunit of the actin regulatory WASH complex, which was lost in fungi (Gomez and Billadeau, 2009), and accordingly, Fam21 proteins were not identified in fungi (fig. 5A). CPI motifs are classified based on the protein, so we generated a fungal Bsp1 consensus sequence based on 33 orthologs to compare to the Bsp1 family CPI motif (fig. 5B and Supplemental Text). The Bsp1 consensus CPI motif was compared against CPI motifs in CARMIL1, CARMIL2, CARMIL3, CIN85, CD2AP, CapZIP, Fam21 and CKIP proteins (fig. 1A) (McConnel et al., 2020). Our sequence analysis revealed that the CPI motif of Bsp1 is closest to those of CKIP and Fam21, which points to a pro-capping function of Bsp1. Taken together these data support three possible origins for Bsp1 proteins: (1) a novel innovation of a fungal CPI motif, (2) a CARMIL divergence that may share an ancestry with the *F. alba* CARMIL tail, or (3) a repurposing of Fam21 after loss of the WASH complex.

The actin capping regulation machinery of *S. cerevisiae*, appears to have diverged significantly from animals and includes: Tda2, ascomycete-specific Aim21, saccharomycotina-specific Aim3 and dikarya-specific Bsp1 proteins (Figure 5A) (Michelot et al., 2013; Shin et al., 2018; Lamb et al., 2021). The innovation of Aim3, an actin capping protein, correlates with loss of ADF/Gelsolin domains in Group II Bsp1s (Figure 5A) and may be functionally connected, however gelsolins and Aim3 capping functions are mechanistically different (Michelot et al., 2013; Nag et al., 2013). CP regulation has diverged significantly, while numerous other proteins that regulate actin dynamics are tightly constrained such as profilin, cofilin, Aip1, twinfilin, CAP/Srv2, coronins, the Arp2/3 complex, formins and tropomyosins. What allows for a large plasticity and specialization in the capping regulation across species and how it relates to the diverse actin functions remains unknown. Bsp1 represents a fungal variation of a pro-capping factor analogous to mammalian CPI motif proteins, which may adapt the actin cytoskeleton to fungal cell biology.

## Author contributions

C.P. Toret had the project idea and performed the phylogenetic analysis. D.R. Hummel performed the cell experiments and analyzed the data. M. Hakala performed the biochemical experiments and analyzed the data. C.P. Toret, D.R. Hummel, M. Hakala and M. Kaksonen designed the experiments and wrote the manuscript.

## Acknowledgments

We are thankful to all the members of the Kaksonen laboratory for technical advice and discussions. We thank Alphée Michelot (IBDM) for the capping protein construct and discussion. We acknowledge Leonardo Scapozza (University of Geneva) for providing an instrument for fluorescence anisotropy assays. This work was supported by the EMBO Postdoctoral fellowship for MH (ALTF 703-2020), and by the Swiss National Science Foundation grant to MK (310030_212288).

## Material & Methods

### Strains, media and plasmids

Strains were created via homologous recombination with PCR cassettes (Janke et al., 2004), and by mating and sporulation. GFP-tagging, deletions and truncations were confirmed by PCR. All fluorescently tagged genes are expressed endogenously. All plasmids and strains used in this study are listed in table S1 and S2, respectively.

### Live cell imaging and analysis

#### Sample preparation

Yeast cells were grown to OD600 between 0.3 and 0.8 at 24 °C with shaking in Synthetic Complete medium without L-Tryptophan (SC-TRP). 40 µl of cells were added to a coverslip coated with Concavalin A. After 5 to 10 minutes of incubation, cells were washed with fresh SC-TRP medium, 40 µl of fresh SC-TRP medium was added and the cells were transferred to the microscope. For DMSO or CK666 treatment, cells were instead washed with SC-TRP medium containing 0.1 mM DMSO or CK666, and 40 µl of DMSO or CK666 medium was added. After 15 minutes of incubation, cells were washed again with DMSO or CK666 medium, 40 µl of DMSO or CK666 medium was added and the cells were transferred to the microscope. For Phalloidin staining, log phase cultures were fixed for 30 minutes in 3.8% formaldehyde, washed and permeabilized in 5% glycine, 0.1% Triton X-100, 1x PBS. We mixed WT and mutant cells with the same mating type, with WT cells expressing Pil1-mCherry, an eisosome marker unrelated to actin processes (Grossmann et al., 2008; Brach et al., 2011). Cells were labeled with Alexa 488 - phalloidin (50 μg/ml in PBS) (Thermofischer) and washed with 1x PBS.

#### Imaging set-up

Epi-fluorescence microscopy images were acquired using a Olympus wide-field IX81 microscope, equipped with a 100x NA1.45 objective and an ORCA-ER CCD camera (Hamamatsu). As illumination source an X-CITE 120 PC (EXFO) metal halide lamp was used. The excitation and emission light when imaging GFP-tagged proteins were filtered through an U-MGFPHQ filter set (Olympus).

#### Image analysis

Images were processed in ImageJ (Schneider et al., 2012). Images were corrected for background fluorescence using the rolling ball algorithm of ImageJ; movies were in addition corrected for photobleaching using the simple ratio algorithm of ImageJ. For particle detection and tracking, the Particle Tracker (Sbalzarini and Koumoutsakos, 2005) of the MosaicSuite was used. Plots were generated using Python 3 controlled by the Jupyter Notebook (Von Rossum & Drake, 2009; Kluyver et al., 2016).

For quantification of Bsp1-GFP and phalloidin patches, patch intensities from z-stack projections were measured and mean with standard deviation of patch intensities was calculated. For phalloidin staining, cells with Pil1-mCherry signal were considered WT in the analysis. Quantification of relative intensities of actin patch components was done as previously described for Bzz1-GFP patches (Hummel and Kaksonen, 2023). Average trajectories of Pan1-GFP patches were obtained as previously described (Picco et al., 2015). Quantification of Cap1-GFP and Bsp1-GFP ring formation was done by manual counting.

### Phylogenetic and sequence analysis

Orthologies were mined with PSI-BLAST and blastp at https://www.ncbi.nlm.nih.gov/. For Bsp1 orthologs, which have low sequence orthology, reciprocal BLAST searches were done first within Saccharomycotina to identify several diverse orthologs, which were used to identify out Ascomycotina orthologs and then expanded out to other clades. The Bsp1 CPI motif consensus site was generated with Jalview.

### Plasmids and protein purification

A BSP1 gene was cloned to pCoofy3 vector (a gift from Sabina Suppmann; Addgene plasmid 43983) as described previously (Scholz et al., 2013). Briefly, BSP1 was amplified from genomic DNA with primers 5’-ctggaagttctgttccaggggcccatgacaaaatatgagcgtgaccctg and 5’-ccccagaacatcaggttaatggcgttacacgcgtgttggaagttttcttc while pCoofy3 was linearized with primers 5’-cgccattaacctgatgttctgggg and 5’-gggcccctggaacagaacttccag using Q5 High-Fidelity 2X Mastermix (New England Biolabs). After DNA fragments were purified from agarose gel, they were annealed using RecA recombinase (New England Biolabs) and transformed into XL10 Gold competent cells (Agilent). Bsp1^1-407^ was amplified using primers 5’-ctggaagttctgttccaggggcccatgacaaaatatgagcgtgaccctg and 5’-ccccagaacatcaggttaatggcgccgtttcttcaggttattcc, and Bsp1^408-470^ with primers 5’-ctggaagttctgttccaggggcccagtattccagaagcaataaaag and 5’-ccccagaacatcaggttaatggcgccgtttcttcaggttattcc. These fragments were annealed to the pCoogy3 vector as described above.

Bsp1 proteins were expressed overnight at 20°C in BL21 pLysS Gold competent cells (Agilent) using autoinduction LB medium (Formedium). Cells were lysed in 50 mM Tris-HCl, pH 8.0, 500 mM NaCl, 2 mM DTT, 2 mM EDTA, 1% Triton X-100 supplemented with PMSF and cOmplete protease inhibitor cocktail (Roche) by sonication. GST-tagged proteins were bound to glutathione sepharose 4B resin (Cytiva) and subsequently washed several times with the lysis buffer. Full length Bsp1 and Bsp1^1-407^ were cleaved from the GST tag with PreScission protease whereas GST-Bsp1408-470 was eluted from the resin using 30 mM glutatione in the lysis buffer without Triton X-100. All proteins were then cleaned with Superdex 200 size exclusion chromatography column equilibrated with 20 mM Tris-HCl, 500 mM NaCl. 2 mM DTT, 2 mM EDTA. Proteins were concentrated and stored with 10% glycerol at -80°C.

Yeast capping protein plasmid for recombinant protein expression in E.coli, pRSFDuet1-6xHis-Cap1/Cap2 (Alphée Michelot, IBDM). Capping protein was expressed and purified as described previously (Antkowiak et al., 2019). After overexpressing the protein in BL21 pLysS cells in autoinduction LB medium, cells we lysed in 20 mM Hepes pH 7.5, 200 mM KCl, 20 mM Imidazole, 0.1 % Triton X-100, 10% glycerol buffer supplemented with PMSF and protease inhibitor cocktail by sonication. Capping protein was then bound to HisTrap Fast Flow column (Cytiva) and eluted with a gradient of 20-300mM imidazole. The final product was cleaned with Superdex 200 size exclusion chromatography column equilibrated with 20 mM Hepes pH 7.5, 200 mM KCl, 10% glycerol and protein was stored in 50% glycerol at -80°C.

### Actin filament cosedimentation assay

Human platelet actin (Cytoskeleton Inc.) was polymerized in F-actin buffer (5 mM Tris-HCl, pH 8.0, 0.2 mM CaCl_2_, 50 mM KCl, 2 mM MgCl_2_, 1 mM ATP) for two hours in room temperature. An indicated concentration of polymerized actin was mixed with 2 µM final concentration of Bsp1 proteins in the F-actin buffer and incubated for 30 minutes at room temperature. Actin filaments were then spinned down with Beckmann TL-100 ultracentrifuge using TLA 100.1 rotor at 75,000 rpm for 30 minutes. Equal amounts of pellet and supernatant were loaded on SDS-PAGE gels. Amounts of Bsp1 protein in pellets and supernatants were measured with Fiji and final graphs were generated using Prism 9 (Graphpad).

### Fluorescence anisotropy

FITC-labeled Bsp1 CPI motif peptide (FITC-RDETVKETKPLVHPNKNRTRGPRRKLPTRV) was synthesized at Genscript. Fluorescence anisotropy measurements were performed in 20 mM HEPES, pH 7.4, 100 mM KCl buffer. 100 nM of FITC-labeled CPI motif peptide and varying concentration of yeast capping protein were incubated for 15 minutes in room temperature before the fluorescence anisotropy was measured with ClarioStar plate reader (BMG labtech). Anisotropy values were calculated with MARS data analysis software (BMG labtech). Final graph was generated with Prism 9 (Graphpad). Data was fitted using the one-site binding equation:

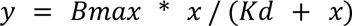

where Bmax is the binding maximum, Kd is the dissociation constant and x is the ligand concentration.

## Data availability

All data generated or analyzed during this study are included in the manuscript and supporting data files.

## Supplemental Figure Legends

**Supplemental figure S1:**
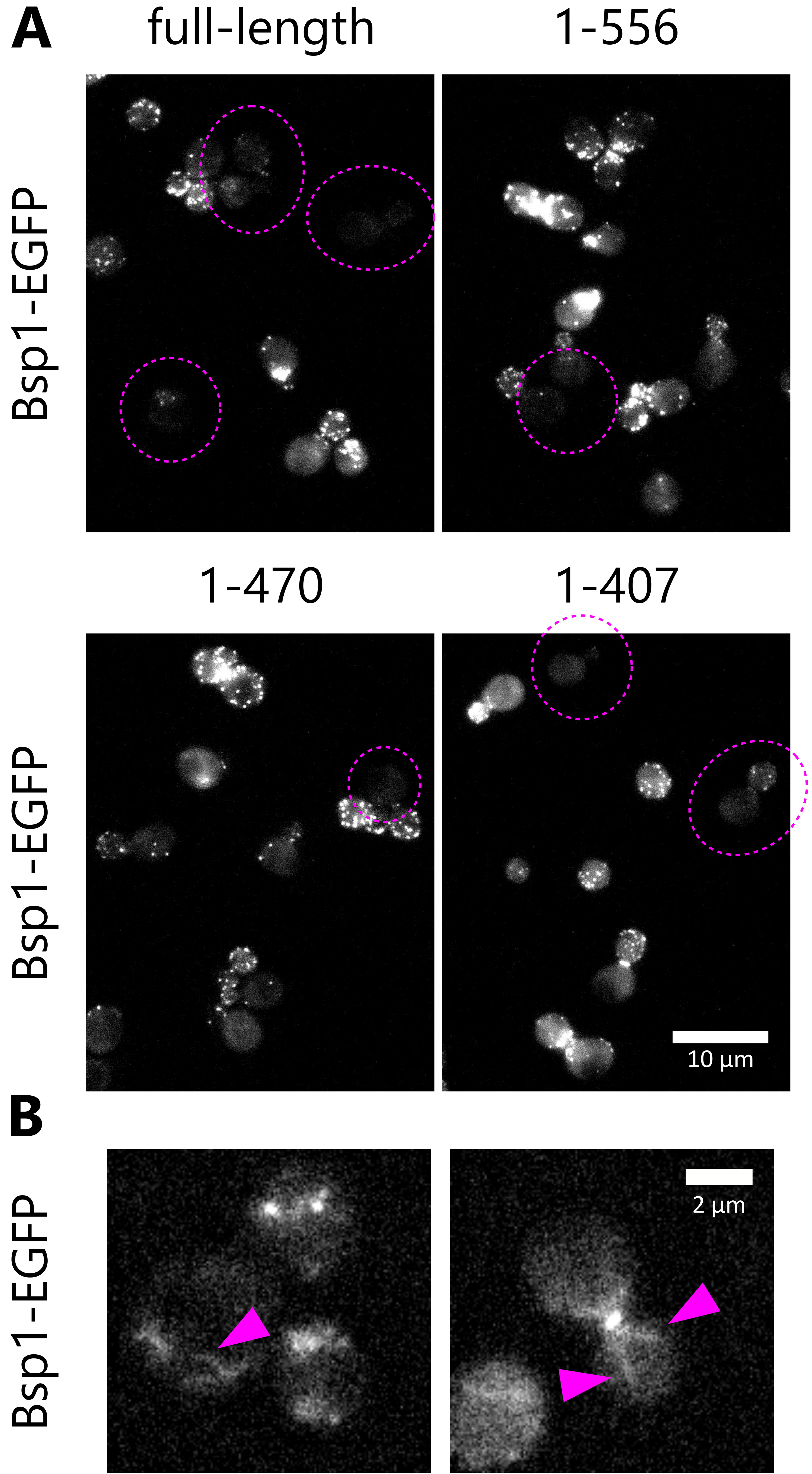
Bsp1 expression heterogeneity. A) Variable Bsp1 expression levels. Example field-of-views of Bsp1-eGFP. Images are projections of z-stacks. Low expression level cells are encircled with a pink dashed line. B) Rare Bsp1-eGFP filamentous localization. Images of representative cells are projections of z-stacks. Pink arrows highlight cable-like structures.

**Supplemental figure S2:**
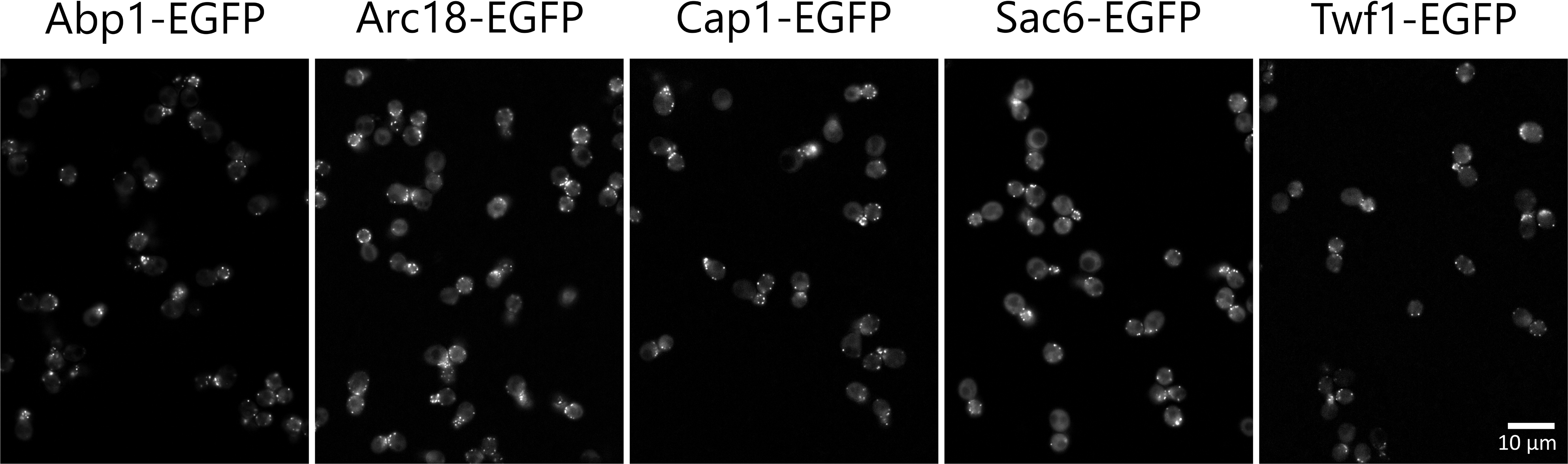
Actin-binding proteins. Example field-of-views of Abp1-eGFP, Arc18-eGFP, Cap1-eGFP, Sac6-eGFP and Twf1-eGFP. Images are single planes and single frames from movies.

## Supplemental Text Legends

**Supplemental Text:** Bsp1 orthologs and CPI motifs

