## Supplemental Table S1 and S2 for "Bsp1, a fungal CPI motif protein, regulates actin filament capping in endocytosis and cytokinesis"

Table S1: Plasmid list

| Name | Alias | Tag | Selection | Source |
| --- | --- | --- | --- | --- |
| pMK0003 | pFA6a-EGFP-HIS6MX | EGFP | hisMX6 | Janke et al., 2004 |
| pMK0086 | pFA6a-KiURA | - | kiURA | Janke et al., 2004 |

Janke C, Magiera MM, Rathfelder N, Taxis C, Reber S, Maekawa H, Moreno-Borchart A, Doenges G, Schwob E, Schiebel E, Knop M. 2004. **A versatile toolbox for PCR-based tagging of yeast genes: new fluorescent proteins, more markers and promoter substitution cassettes**. *Yeast* **21**:947–962. doi:10.1002/YEA.1142

Table S2: Strain list

| Figure | Name | Genotype | Strain number |
| --- | --- | --- | --- |
| 1&4 | Bsp1-eGFP | MATa, his3-Δ200, leu2-3,112, ura3-52, lys2-801, BSP1-EGFP::his3MX6 | MKY4816 |
| 1&4 | Bsp1(CPI Δ)-eGFP | MATa, his3-Δ200, leu2-3,112, ura3-52, lys2-801, BSP1(1-556)-EGFP::his3MX6 | MKY4818 |
| 1 | Bsp1(tail Δ)-eGFP | MATa, his3-Δ200, leu2-3,112, ura3-52, lys2-801, BSP1(1-470)-EGFP::his3MX6 | MKY4896 |
| 1 | Bsp1(WH2 & tail Δ)-eGFP | MATa, his3-Δ200, leu2-3,112, ura3-52, lys2-801, BSP1(1-407)-EGFP::his3MX6 | MKY4897 |
| 3 | Twf1-eGFP | MATα, his3-Δ200, leu2-3,112, ura3-52, lys2-801, TWF1-EGFP::his3MX6 | MKY4747 |
| 3 | Twf1-eGFP *bsp1Δ* | MATα, his3-Δ200, leu2-3,112, ura3-52, lys2-801, TWF1-EGFP::his3MX6, bsp1Δ::klURA3 | MKY4757 |
| 3 | Abp1-eGFP | MATα, his3-Δ200, leu2-3,112, ura3-52, lys2-801, ABP1-EGFP::his3MX6 | MKY2834 |
| 3 | Abp1-eGFP *bsp1Δ* | MATα, his3-Δ200, leu2-3,112, ura3-52, lys2-801, ABP1-EGFP::his3MX6, bsp1Δ::klURA3 | MKY4755 |
| 3 | Sac6-eGFP | MATα, his3-Δ200, leu2-3,112, ura3-52, lys2-801, SAC6-EGFP::his3MX6 | MKY3863 |
| 3 | Sac6-eGFP *bsp1Δ* | MATα, his3-Δ200, leu2-3,112, ura3-52, lys2-801, SAC6-EGFP::his3MX6, bsp1Δ::klURA3 | MKY4753 |
| 3 | Arc18-eGFP | MATa, his3-Δ200, leu2-3,112, ura3-52, lys2-801, ARC18-EGFP::his3MX6 | MKY2150 |
| 3 | Arc18-eGFP *bsp1Δ* | MATa, his3-Δ200, leu2-3,112, ura3-52, lys2-801, ARC18-EGFP::his3MX6, bsp1Δ::klURA3 | MKY4782 |
| 3 | Pan1-eGFP | MATa, his3-Δ200, leu2-3,112, ura3-52, lys2-801, PAN1-EGFP::his3MX6 | MKY0676 |
| 3 | Pan1-eGFP *bsp1Δ* | MATa, his3-Δ200, leu2-3,112, ura3-52, lys2-801, PAN1-EGFP::his3MX6, bsp1Δ::klURA3 | MKY4780 |
| 3 | *bsp1Δ* | MATa, his3-Δ200, leu2-3,112, ura3-52, lys2-801, bsp1Δ::klURA3 | MKY4746 |
| 3 | Pil1-mCherry | MATa, his3-Δ200, leu2-3,112, ura3-52, lys2-801, PIL1-MCHERRY::KANMX4 | MKY0144 |
| 3&4 | Cap1-eGFP | MATa, his3-Δ200, leu2-3,112, ura3-52, lys2-801, CAP1-EGFP::his3MX6 | MKY2865 |
| 4 | Cap1-eGFP | MATα, his3-Δ200, leu2-3,112, ura3-52, lys2-801, CAP1-EGFP::his3MX6 | MKY3326 |
| 3&4 | Cap1-eGFP *bsp1Δ* | MATα, his3-Δ200, leu2-3,112, ura3-52, lys2-801, CAP1-EGFP::his3MX6, bsp1Δ::klURA3 | MKY4775 |
| 4 | Cap1-eGFP *bsp1(CPIΔ)* | MATα, his3-Δ200, leu2-3,112, ura3-52, lys2-801, CAP1-EGFP::his3MX6, bsp1(1-556)::klURA3 | MKY4940 |
| - | *bsp1Δ* | MATα, his3-Δ200, leu2-3,112, ura3-52, lys2-801, bsp1Δ::klURA3 | MKY4758 |
| - | *bsp1(CPIΔ)* | MATa, his3-Δ200, leu2-3,112, ura3-52, lys2-801, bsp1(1-556)::klURA3 | MKY4921 |
